# Zinc nano-fertilization enhances wheat productivity and biofortification

**DOI:** 10.1101/2023.01.06.522993

**Authors:** Achchhelal Yadav, Pramila Krishnan, Monika Kundu, Ram Swaroop Bana, Shaloo, Anil K. Choudhary, Y.S. Shivay, Samrath Lal Meena, Shbana Begam, Samarth Godara, Rajeev Ranjan, Sudhir Kumar, Sunita Yadav, M.S. Nain, Teekam Singh, Abhijeet Yadav, Rishi Raj

## Abstract

Zinc (Zn) malnutrition has emerged as one of the major health challenges in developing nations across the globe. Development of Zn management protocols in staple food crops using modern scientific tools to enhance Zn concentration in grains along with augmented crop yields became utmost necessary. In this context a 2-year experiment was carried out to assess the effects of zinc oxide nanoparticles (ZnO-NPs) vis-à-vis bulk zinc sulfate (ZnSO_4_) on wheat growth, yield and Zn concentration in plant parts. Four levels of application of ZnO-NPs (0, 20, 25 and 30 mg kg^-1^) were compared with ZnSO_4_ (equal to Zinc concentration in ZnO-NPs). Results revealed that seed vigor was significantly (p <0.05) higher under 25 and 30 mg kg^-1^ soil ZnO-NPs treatments over ZnSO_4_. Among the crop yield parameters such as tillers (plant^-1^), grain weight (plant^-1^), biomass (plant^-1^) and grain yield were significantly (p <0.05) higher under ZnO-NPs 25 mg kg^-1^ treated soil as compared to any other treatment. Zinc concentration in grains increased with dose of ZnO-NPs and it was significantly more than ZnSO_4_ treated soil at each treatment level. ZnO-NPs and ZnSO_4_ treatments did not affect photosynthetic rate and chlorophyll (SPAD) content significantly. In conclusion, 25 mg kg^-1^ ZnO-NPs application could be recommended in wheat cultivation to improve growth, yield and grain Zn biofortification.

## Introduction

In the modern era nanotechnology is being used in almost all existing science fields [1–3]. As per the definition of nanotechnology, a particle should have a dimension in range of 1 to 100 nanometers [4]. The size of material in nano-range increases its biocompatibility, surface area, electrical conductivity and antimicrobial activities, which makes it suitable for multi-uses [5-7]. Among the various NPs, zinc oxide (ZnO)-NPs are being used widely and their rank is fourth across the NPs [1]. In the recent past, the application of micronutrient NPs has emerged as a significant part of nutrient management schedules of crop plants owing to their potential advantages on productivity and quality gains [8–10]. Presently, soil and foliar applications of NPs in crops are under practice [11–15].

Zinc is an essential plant micronutrient and an integral part of biochemical processes of plants including protein synthesis [16–18]. The ZnO-NPs can perform the role as fertilizers, pesticides, growth regulators and herbicides [19–20]. The deficiency of micronutrients like Zn in soils reduces crop productivity as well as grain nutritional quality [21–23]. The constant decline in levels of soil Zn content across diverse agro-ecologies, has posed an alarming challenge to improve Zn content of grain and therefore Zn management has emerged as most important area of research during recent period [24]. The high solubility, small molecular weight and lower thermal stability of conventional fertilizers result in reduced efficacy and enhanced environmental pollution [25–27].

On the contrary, Zn application as nano-formulations enhances the fertilizer use efficiency due to greater surface area and specific properties. Further, lower runoff and leaching losses of nanoparticles would reduce soil and air pollution, while providing higher crop yields and nutrient uptake [28, 29]. Further, application of Zn in various formulations, including NPs, assists in enrichment of edible plant parts in Zn, resulting in reduced hidden hunger and Zn malnutrition [30, 31].

Zinc is also a crucial trace element for human health that has numerous regulatory and important enzymatic activities involved in various metabolic pathways and biochemical functions. After iron, Zn is the most profuse microelement available in the human body [32]. In the developing world, one third population is suffering with the deficiency of Zn, especially children and pregnant women [33–35]. Paucity of Zn in human body causes suboptimal health status, increased morbidity, mortality, physiological alterations, shortfall in growth, and infectious diseases [36]. The poor dietary variation and low intake of Zn may be the reasons for Zn deficiency in the developing nations [36, 37]. Wheat is an important cereal that provides required calories and proteins across the world. The production and consumption of Zn-enriched cereals, such as wheat, could be one of the most powerful weapons for fighting against Zn malnutrition. The ZnO-NPs are being used in industry for several decades. However, their application in the agriculture are not potentially explored. So far no much information is available on effect of soil application of ZnO-NPs on agronomic biofortification of wheat. Further, numerous focused information is available on productivity enhancement due to ZnO-NPs foliar application, however, detailed information on effects of ZnO-NPs on crop growth, yield attributing characters, rooting behavior and physiological plant parameters is not available. Likewise, knowledge gap also exists on Zn partitioning behaviour in different wheat plant parts under diverse Zn nutrition systems, including soil application of ZnO-NPs. Therefore, the present study was undertaken to evaluate the effect of ZnO-NPs on wheat growth, yield and physiological processes in the plant system. Furthermore, the current work also aimed to understand Zn partitioning in different plant parts under ZnO-NPs and under ZnSO_4_. We hypothesized that ZnO-NPs application in soil may result in enhancement of wheat growth, yield and Zn biofortification.

## Method and materials

### Description of site

Present study was undertaken during 2019-20 and 2020-21 winter seasons at ICAR-Indian Agricultural Research Institute, New Delhi, India, in plastic pots (top diameter 32 and 34 cm height). In the experiment, seven treatments were evaluated with six replications in the completely randomized design (Table 2). The soil for pot filling (0-15 cm soil profile) was digged and collected from MB 4C research farm of the Institute. The sand, silt and clay percent of digged soil were 64.0, 16.8 and 19.2%, respectively. The texture of soil was sandy loam [38]. The soil was dried, sieved with 1.0 cm sieve and properly mixed and filled in pots @ 17 kg pot^-1^. The soil had electrical conductivity (1:2 soil: water) of 0.15 dSm^-1^, pH 7.7, zinc concentration 0.78 mg kg^-1^ and carbon content of 0.51% at the time of sowing. Experimental site is located in the Indo-Ganagetic plains in north-west India at 28° 38’ 23” N, 77° 09’27” E at an altitude of 228.6 m above mean sea level. It belongs to sub-tropical semiarid region and had total of rainfalls 306 and 71.1 mm from November 2019 to April 2020 and from November 2020 to April 2021 (wheat growing periods), respectively. The mean maximum temperatures were 24.7 and 11.4 and minimum temperature were 11.4 and 10.6 °C from November 2019 to April 2020 and November 2021 to April 2021, respectively. The respective mean relative humidity during both the study periods were 67.7 and 62.7%.

**Table 1.** Description of treatments.

| Symbol | Pot soil treatment |
| --- | --- |
| T1 | Control |
| T2 | Application of ZnO NPs in pot soil 20 mg kg<sup>-1</sup> |
| T3 | Application of ZnO NPs in pot soil 25 mg kg<sup>-1</sup> |
| T4 | Application of ZnO NPs in pot soil 30 mg kg<sup>-1</sup> |
| T5 | Application of ZnSO<sub>4</sub> in pot soil 49.2 mg kg<sup>-1</sup> (Zn equivalent to T2) |
| T6 | Application of ZnSO<sub>4</sub> in pot soil 61.5 mg kg<sup>-1</sup> (Zn equivalent to T3) |
| T7 | Application of ZnSO<sub>4</sub> in pot soil 73.8 mg kg<sup>-1</sup> (Zn equivalent to T4) |

**Table 2.** The effect of ZnO-NPs and ZnSO_4_ on plant growth parameters. Two-year data of seedling vigor I, photosynthetic rates, SPAD values and tillers per plant are given. Mean data of 2 years mentioned Parameters crop grown under ZnO-NPs as well as ZnSO_4_ treatment are compared with the control treatment (T1). Data of seedling Vigor I, and tiller (plant-1) shown in bold are significantly different over control treatment data.

| Plant Growth | Seedling Vigor I | | | Photosynthetic rate<br>($\mu\text{mol m}^{-2} \text{s}^{-1}$) | | | SPAD value | | Tillers per plant | | | |
| --- | --- | --- | --- | --- | --- | --- | --- | --- | --- | --- | --- | --- |
|  | 2019-20 | 2020-21 | Mean | 2019-20 | 2020-21 | Mean | 2019-20 | 2020-21 | Mean | 2019-20 | 2020-21 | Mean |
| T1 | 1998 | 2294 | 2146 | 9.9 | 10.3 | 10.1 | 51.1 | 50.5 | 50.8 | 6.4 | 6.5 | 6.5 |
| T2 | 2109 | 1981 | 2045 | 8.8 | 11.2 | 10 | 54.7 | 56 | 55.3 | 6.9 | 7 | 7 |
| T3 | 2087 | 2225 | 2156 | 11.3 | 13.5 | 12.4 | 54.5 | 52 | 53.2 | **8.2** | **8** | **8.1** |
| T4 | **2132** | 1964 | 2048 | 9.6 | 13 | 11.3 | 55.1 | 51 | 53 | 6.5 | 6.7 | 6.6 |
| T5 | 1865 | 2067 | 1966 | 8.8 | 8.21 | 8.5 | 49 | 53 | 51 | 6.8 | 6.5 | 6.7 |
| T6 | 1856 | 1943 | **1899** | 8.9 | 10.2 | 9.5 | 53.3 | 51 | 52.2 | 6.8 | 6.3 | 6.6 |
| T7 | 1788 | 1632 | **1710** | 10.8 | 13.3 | 12 | 51 | 47 | 49 | 6.4 | 6.2 | 6.3 |
| CD (p=0.05) | 247.7 | 206.9 | 232.1 | ns | ns | ns | ns | ns | ns | 1.66 | 1.50 | 1.46 |
| SE(m) | 67.8 | 82.7 | 79.7 | ns | ns | ns | ns | ns | ns | 0.57 | 0.46 | 0.51 |

ns-non significant

### Procurement of ZnO-NPs and ZnSO_4_

The ZnO-NPs (Type I) approximately 30 nm size were procured from Sisco Research Laboratories (SRL) Pvt. Ltd. (Batch No. 8628965; Manufactured date 06 June, 2017; Expiry/Retest date 6^th^ June 2022). The ZnO-NPs were characterized using transmission electron microscopy (Fig 1) [39]. The ZnSO_4_ was also purchased from SRL (Batch No.:011219).

**Fig. 1.**
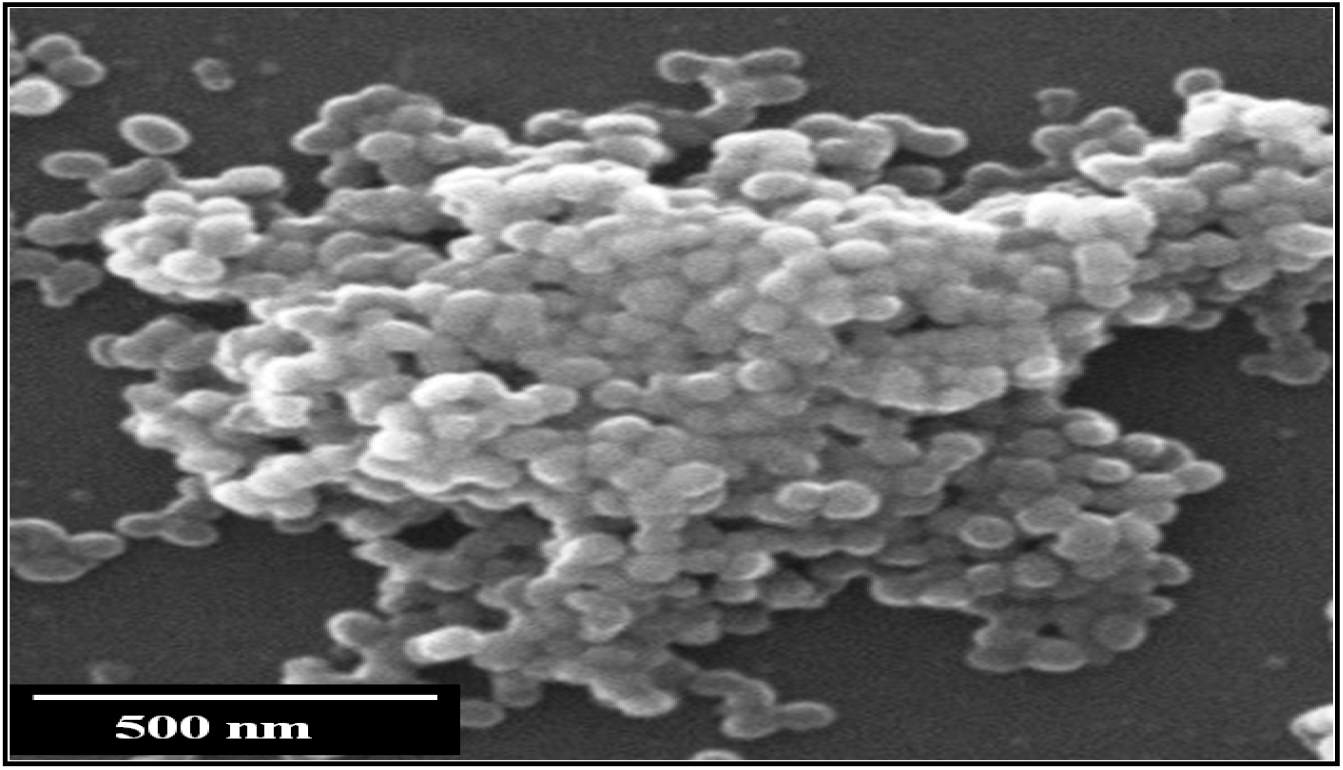
Transmission microscopy picture of ZnO-NPs.

### Experiment on Germination

The seeds of wheat were selected carefully and were sterilized for 10 minutes with solution of 1% hydrogen peroxide (H_2_O_2_) and washed 3 times with double distilled water for insuring their sterility [40]. Solutions of different concentration of ZnO-NPs (0, 340, 425, 510 mg litre^-1^) and 836, 1050 and 1254 mg litre^-1^ ZnSO_4_ (equivalent to ZnO-NPs) were prepared by dissolving ZnO-NPs and ZnSO_4_ in double distilled water (volume was maintained 1 litre). Concentrations of ZnO-NPs and ZnSO_4_ dispersions were equivalent to the total concentrations which were applied to the pot soil during crop growth period. Thirty seeds of wheat were put in germination paper to assess germination (%) and seed vigor. A 30 ml solution of different concentration of ZnO-NPs and bulk ZnSO_4_ were poured in each germination paper in six replicates. The germination papers were kept in a dark incubator at 29 ±1.1 °C. Germination of seeds was recorded after every 24 hours till 7 days. Seedlings length, seedlings biomass and root biomass were determined after 7 days, the samples were kept in the oven (at 65 °C) for drying. Pot soil was washed and roots samples were collected in triplicate for the measurement of root parameters.

### Crop fertilization

The optimum amount of N (half dose; 6.5 g pot^-1^), P_2_O_5_ (3.8 g pot^-1^) and K_2_O (3.1 g pot^-1^) fertilizers were applied in the pot soil during the sowing. The 50% of nitrogen dose was given at the time of sowing and remaining doses were applied in two equal parts at vegetative and flowering stages, respectively. The full doses of P_2_O_5_ and K_2_O were applied at once during the sowing. Apart from required N, P and K fertilizers, the pot soil was treated with ZnO-NPs as well zinc sulfate (ZnSO_4_.7H_2_O). The ZnO-NPs and ZnSO_4_ were weighed using an electronic balance (Aair Dutt ViBRA AJ). The ZnO-NPs as well bulk ZnSO_4_ solution were prepared with double distilled water. Stirrer was used to mix properly ZnO-NPs and bulk ZnSO_4_ in double distilled water. This solution was poured in the pot soils in six different treatments [T1: 0 mg kg^-1^; T2: 20 mg kg^-1^ ZnO-NPs; T3: 25 mg kg^-1^ ZnO-NPs; T4: 30 mg kg^-1^ ZnO-NPs; T5: 20 mg kg^-1^ ZnSO_4_ (equivalent to Zn Content in T2); T6: 25 mg kg^-1^ ZnSO_4_ (equivalent to Zn in T3); and T7: 30 mg kg^-1^ ZnSO_4_ (Zn equivalent to T4)] before the sowing and mixed appropriately (Table 1). The equivalent amount of Sulphur of ZnSO_4_ was applied in the ZnO-NP treatments using elemental Sulphur. The sowing of crop was done at 5 cm plant to plant distance and 22.5 cm row to row distance during both the cropping seasons. The sowing of wheat in pot soil was done on 20 November 2019 and 18 November 2020 with 2-rows of wheat in each pot. The harvesting of crops was done on 17 April and 14 April of 2020 and 2021, respectively

### Measurement of plant growth parameters

The seedling vigor index 1 after 7 days of planting was determined using the formula germination (%) × shoot length of the seedling. The photosynthetic rate (Pn), stomatal conductance chlorophyll content was determined at the pre-flowering stage of crop growth. The photosynthetic rate was measured using LI-6400XT Portable Photosynthetic System (Model No. LI-6400XT, LI-COR, USA). The chlorophyll content was determined using the SPAD (SPAD-502 Plus).

### Determination of yield and yield attributes

Five representative plants were selected from each treatment and harvested at maturity. Later, plant samples were separated into stem, leaves and spikes and exposed to heat burst in microwave. These samples were dried in oven at 65 °C for 48 hours [41]. The crop yield and yield attributes such as spike length (cm), spikelets (spike^-1^), number of grains (spike^-1^), grain weight (g spike^-1^), biomass (g plant^-1^), grain yield (g plant^-1^), harvest index (grain yield divided by biomass multiplied by 100) and 1,000-grain weights (g) were determined.

### Determination of zinc in different parts of wheat

The 0.5 g fine grinded wheat samples of root, stem, leaves and grains were taken and digested with the mixed solution of concentrated HNO_3_ and HClO_4_ acids (in 9:4 ratio) in a conical flask [42]. The flasks were kept for digestion on hot plate for heating (3.5 hours) or until a colorless residue was left in the digestion vessel. After cooling, the residue was mixed with a diluted solution of H_2_SO_4_ (0.1 N) and volume was made up to 100 ml. The digested samples were taken for estimating Zn using atomic absorption spectrometry at 213.9 nm (AAS PLUS, Motras Scientific, India).

### Statistical analysis

The crop growth, yield and quality attributes of wheat (variety: HD 2967) were assessed using 9.3 SAS statistical software. One-way analysis of variance (ANOVA) was used to evaluate the effects of ZnO-NPs and their counterpart conventional fertilizer ZnSO_4_ application on soils. The data were compared at 95 %confidence level (p < 0.05) to test statistical significance.

## Results

### Effect of ZnO-NPs on seedling shoot lengths, biomass and seedling vigor I

Among different zinc application treatments (through zinc oxide nanoparticles and zinc sulfate forms), the germination was higher in the initial four days under each dose of ZnO-NPs (T2-T4) vis-à-vis in ZnSO_4_ (T5-T7), but it was non-significantly different across the treatments (T1-T7) (data not given). The seedling growth was assessed using shoot length, root length and dry weight of seedlings (Fig 2). Results showed that the root length (cm) on the seventh day after sowing was significantly more under 20 and 25 mg litre^-1^ ZnO solution (21.3% in T2 and 6.79% in T3) treatment, however it reduced notably (−7.8 to −32.0%) under other treatments of ZnO-NPs as well as ZnSO_4_ (T4-T7) as compared to control (Fig 3A). The shoot length relative (%) to control was significantly more (p <0.05) under T2 (15.8%) and T3 (5.0%) treatments, while no significant changes were observed in other treatments (Fig 3B). Seedling dry weight showed non-significant enhancement under the ZnO-NPs treatments, however, the ZnSO_4_ treatments resulted in a significant reduction over the control (Fig 3C).

**Fig. 2.**
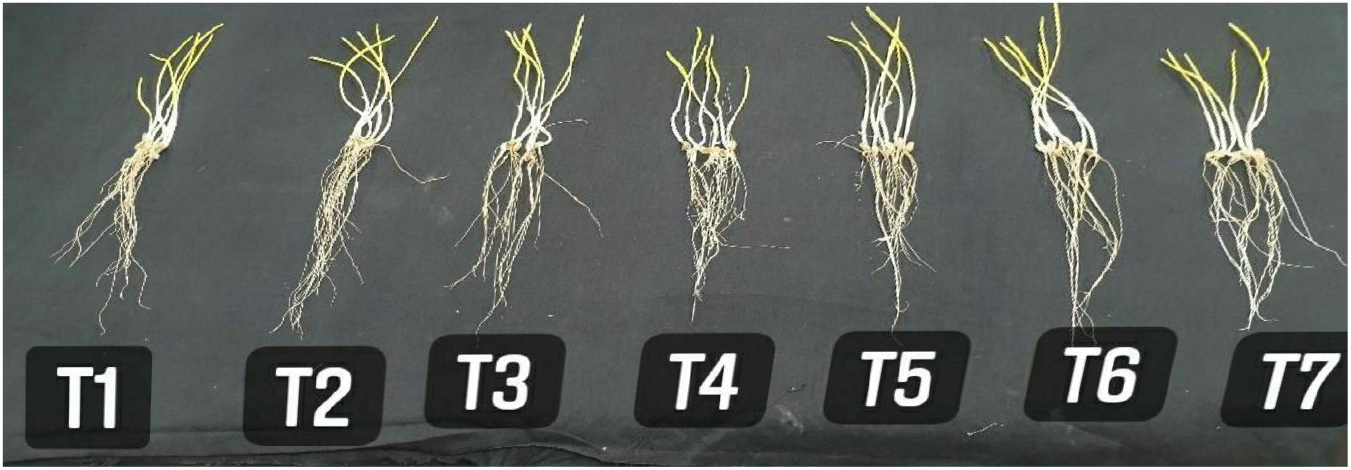
Seedlings grown under different in ZnO and ZnSO_4_ treatments.

**Fig. 3.**
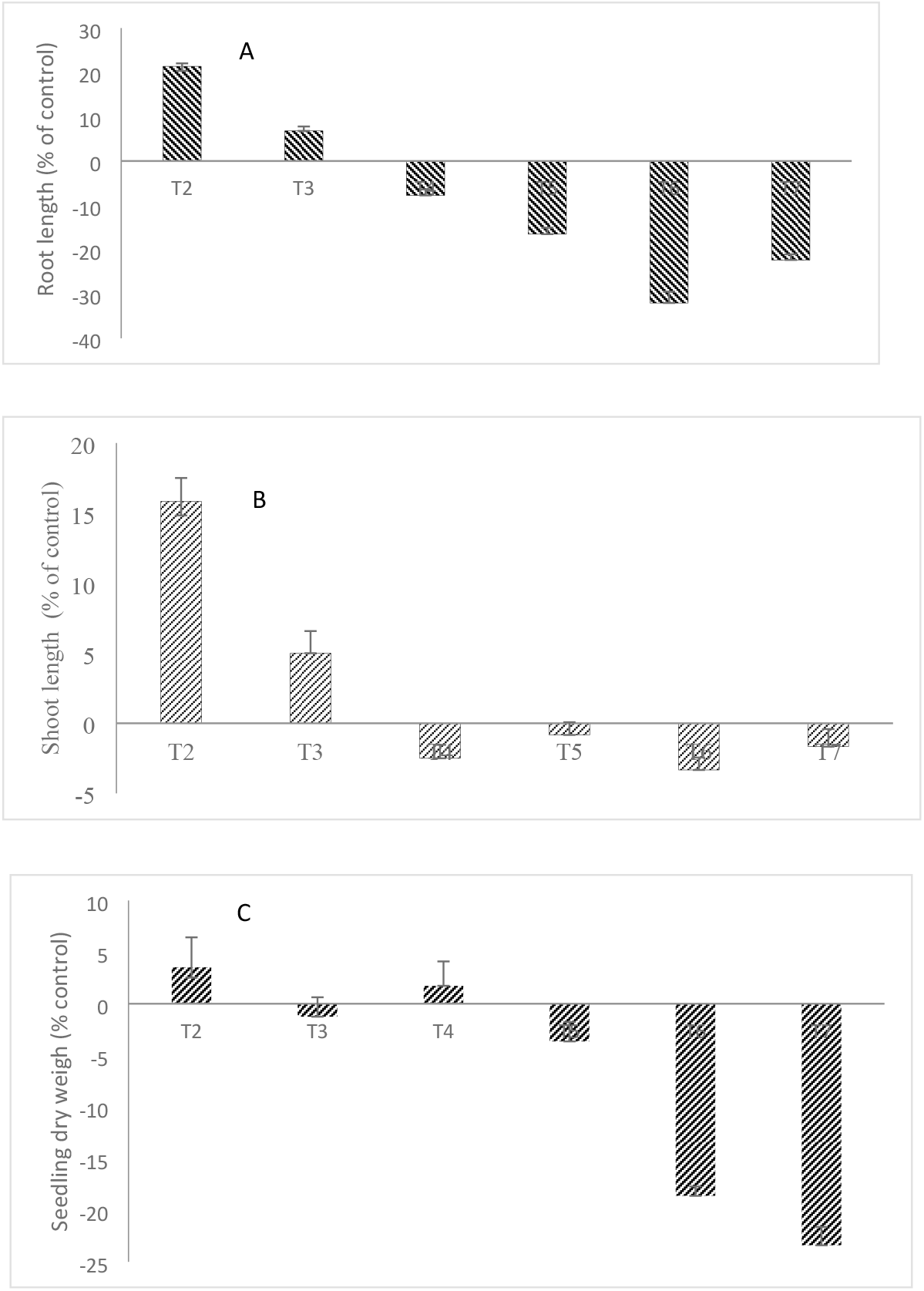
Effect of ZnO-NPs and ZnSO_4_ on (A) root length (B) shoot length and (C) seedling dry weight. The comparison of seedling dry was done with the dry weight of seedling of treatment T1 and the percentage changes are reported.

### Effect of ZnO-NPs and ZnSO_4_ on photosynthetic rate and SPAD

The photosynthetic rate and SPAD values did not change significantly under ZnO-NPs and under ZnSO_4_ treatments over the control (Table 2). Tiller plant^-1^ (ranged from 6.2 to 8.2) were recorded 24 % higher under the T3 treatment (ZnO-NPs 25 mg kg^-1^) over the control (T1) (Table 2). Similarly, as compared to T6, a 22.7% enhancement in the number of tillers was noticed under T3.

### Effect of ZnO-NPs on yield attributes

The mean of two years of biomass (plant^-1^) ranged from 11.9 g (plant^-1^) in control and 14.5 and 12.8 g plant^-1^ in ZnO-NPs and ZnSO_4_ treatments, respectively (Table 3). Data revealed that the biomass (plant^-1^) was significantly higher under 25 mg kg^-1^ ZnO-NPs treated soil (T3) than that of any other treatment of ZnO-NPs and ZnSO_4_. The percent biomass increase under T3 was 22.8% as compared to control. The comparison of biomass between T3 and T6 treatments showed that 25 mg kg^-1^ (T3) treated soil recorded 13.3% higher biomass (plant^-1^) over ZnSO_4_ treated soil (T6), which suggest that the ZnO-NPs use is more efficient that than its counterpart ZnSO_4_ fertilizer (Table 3). The grains (spike^-1^) were statistically non-significant (p>0.05) across the treatments. Grain weight (g) ranged from 1.1-1.3 g spike^-1^. The grain weight (g spike^-1^) was significantly higher under ZnO-NPs treatment than any other treatment. It increased by 18.2% under T3 than T6 (Table 3). Similar to biomass, grain yield (g plant^-1^) ranged from 5.9 g plant^-1^ (control) to 7.8 g plant^-1^ (ZnO-NPs). Data revealed that the grain yield was significantly (p<0.05) higher in T3 than that of any other treatment (Table 3). Comparative assessment between grain yields of T3 and T6 treatments indicated that the grain yield under T3 was 16.6% greater than T6 (Table 3). This suggests that 25 mg kg^-1^ ZnO-NPs treatment (T3) remained more efficient than its equivalent ZnSO_4_ (T6) treatment for grain yield of wheat. The 1,000-grain weight and harvest index (%) did not change significantly across the treatments (Table 3).

**Table 3.** The effect of ZnO-NPs and ZnSO_4_ on yield parameters of wheat. Two-year data of biomass, grain weight (g-plant^-1^), number of grains (spike^-1^) grain yield (g-plant^-1^), 1000 grain weight (g) and harvest (%) are given. Mean of 2-year data of mentioned parameters crop grown under ZnO-NPs as well as ZnSO_4_ treatment are compared with the control treatment (T1). Data of biomass (g-plant^-1^), grain weight (g-spike^-1^) and grain yield (g-plant^-1^) shown in bold are significantly different from control treatment and themselves.

| Treatment | Biomass (g-plant<sup>-1</sup>) |  |  | Grain weight (g-spike<sup>-1</sup>) |  |  | Number of grains (spike<sup>-1</sup>) |  |  | Grain yield (g-plant<sup>-1</sup>) |  |  | 1,000 grain weight (g) |  |  | Harvest Index (%) |  |  |
| --- | --- | --- | --- | --- | --- | --- | --- | --- | --- | --- | --- | --- | --- | --- | --- | --- | --- | --- |
|  | 2019-20 | 2020-21 | Mean | 2020-21 | 2020-21 | Mean | 2019-2020 | 2020-21 | Mean | 2019-2020 | 2020-21 | Mean | 2019-2020 | 2020-21 | Mean | 2019-2020 | 2020-21 | Mean |
| T1 | 11.2 | 12.3 | 11.8 | 1 | 1.2 | 1.1 | 37.0 | 41.4 | 39.3 | 5.7 | 6.0 | 5.9 | 27.0 | 29 | 28 | 50.9 | 48.8 | 49.8 |
| T2 | **12.1** | **13.5** | **12.8** | 1.1 | **1.3** | **1.2** | (g) | 46.4 | 41.8 | 6.3 | 6.5 | 6.4 | 29.4 | 28 | 28.7 | 52.1 | 48.1 | 50.0 |
| T3 | **14.7** | **14.3** | **14.5** | 1.3 | **1.3** | **1.3** | 42.6 | 44.4 | 43.5 | **8.1** | 7.4 | **7.8** | 30.5 | 29.3 | 29.9 | 55.1 | 51.7 | 53.4 |
| T4 | **13.1** | **12.8** | **13.0** | 1 | 1 | 1 | 34.5 | 33.3 | 33.4 | 6.7 | 7.0 | 6.9 | 29.8 | 30 | 29.9 | 51.1 | 54.7 | 52.9 |
| T5 | 11.7 | 12.3 | 12.0 | 0.9 | 1 | 0.9 | 30.0 | 32.9 | 29.7 | 6.1 | 6.0 | 6.1 | 30.1 | 30.4 | 30.3 | 49.2 | 45.5 | 47.3 |
| T6 | **12.4** | **13.2** | **12.8** | 1 | 1.1 | 1.1 | 34.0 | 37.0 | 37.2 | 6.3 | 7.0 | 6.7 | 29.4 | 29.7 | 29.6 | 53.8 | 56.9 | 55.4 |
| T7 | **12.2** | **11.6** | 11.9 | 1 | 1 | 1 | 33.7 | 33.3 | 33.4 | 6.5 | 6.6 | 6.6 | 29.7 | 30 | 29.9 | 53.3 | 56.9 | 55.0 |
| CD (p=0.05) | 0.44 | 0.43 | 0.48 | 0.219 | 0.199 | 0.139 | ns | ns | ns | 1.83 | 1.87 | 1.85 | ns | ns | ns | ns | ns | ns |
| SE(m) | 0.153 | 0.157 | 0.161 | 0.078 | 0.092 | 0.082 | ns | ns | ns | 0.625 | 0.678 | 0.710 | ns | ns | ns | ns | ns | ns |

ns-non significant

### Effect of ZnO-NPs on Zn content in below- and above-ground biomass

The Zn concentration in roots increased from 31.7 (T1) to 78.9 mg kg^-1^(T4). Under the ZnO-NP treatments it ranged from 41.9 to 78.9 mg kg^-1^ (T2-T4) and in ZnSO_4_ soil treatments it ranged from 32.2 to 37.2 mg kg^-1^ (T5-T7) (Table 4). Concentration of Zn in roots of T4 was 2.48 times more than that of T1 (Table 4). Similarly, the concentration of Zn in roots under ZnSO_4_ treated soil (T7) was higher by 17.4% over the control treatment (T1). Comparison of data between treatments T4 and T7 (highest doses of ZnO NPs and ZnSO_4_) also suggests that the concentration of Zn in roots under T4 (counter part of T7) was significantly more (112.1%) as compared to T7 treatment (Table 4). The Zn concentration in stem ranged from 20.2 mg kg^-1^ (control) to 30.7 and 27.2 mg kg^-1^ under ZnO-NPs and ZnSO_4_ treated soils respectively. Zn concentration was more by 51.9% and 34.6% at T4 and T7 soil treatment over the T1 (control). Concentration of zinc was increased by 12.9% under T4 over T7. From the above, it could be concluded that the Zn absorption was taking place more efficiently under ZnO-NPs treated soil than ZnSO_4_ applied soil (Table 4).

**Table 4.** The zinc concentration (mg kg^-1^) of root, stem, leaf and grains of wheat. Two-year data of zinc concentration in root, stem, leaf and grains (%) are given. Mean data of Zn contents in crop plants under ZnO-NPs as well as ZnSO_4_ treatment are compared with the control treatment (T1) parameter. Data of Zn concentrations in different plant part shown in bold are significantly (p<0.05) different over the control (T1) as well themselves.

| Treatment | Root |  |  | Stem |  |  | Leaf |  |  | Grains |  |  |
| --- | --- | --- | --- | --- | --- | --- | --- | --- | --- | --- | --- | --- |
|  | 2019-20 | 2020-21 | Mean | 2019-20 | 2020-21 | Mean | 2019-20 | 2020-21 | Mean | 2019-20 | 2020-21 | Mean |
| T1 | 30.0 | 32.8 | 31.4 | 21.5 | 18.9 | 20.2 | 7.1 | 7.7 | 7.4 | 30.1 | 33.2 | 31.7 |
| T2 | 43.8 | 40.0 | 41.9 | 24.7 | 22.4 | 23.5 | **12.7** | 10.2 | 11.4 | 36.0 | 33.4 | 34.7 |
| T3 | **65.0** | **61.7** | **63.4** | 25.8 | **27.4** | **26.6** | **17.0** | **19.4** | **18.2** | **40** | **42.1** | **41.0** |
| T4 | **77.1** | **80.7** | **78.9** | **32.0** | **29.5** | **30.7** | **20.2** | **26.7** | **23.5** | **50.1** | **48.0** | **49.0** |
| T5 | 32.0 | 34.4 | 33.2 | 20.7 | **24.9** | 22.8 | **22.1** | **20.1** | **21.1** | 33.0 | 35.2 | 34.1 |
| T6 | 34.7 | 35.7 | 35.2 | 22.7 | 23.8 | 23.2 | **23.7** | **24.2** | **23.9** | 34.7 | 35.6 | 35.2 |
| T7 | **36.9** | 38.0 | 37.4 | 26.0 | **28.3** | **27.2** | **28.3** | **26.6** | **27.4** | **39.8** | 33.4 | 36.6 |
| CD (p <0.05) | 6.77 | 7.31 | 7.47 | 5.52 | 5.13 | 4.29 | 4.31 | 3.99 | 4.02 | 7.01 | 6.88 | 6.95 |
| SE(m) | 2.89 | 2.65 | 2.44 | 1.72 | 1.62 | 1.67 | 1.27 | 1.33 | 1.31 | 2.41 | 2.19 | 2.27 |

The Zn concentration in crop leaves was significantly increased with increase in the levels of ZnO-NPs as well as ZnSO_4_ in pot soils. The mean Zn concentration in leaves ranged from 7.4 mg kg^-1^(control) to 23.5 and 27.4 mg kg^-1^ in ZnO-NPs and ZnSO_4_ treated soils, respectively. It is obvious from data that the mean of Zn concentration in leaves was more (36.4%) under ZnSO_4_ treated soil over ZnO-NPs treated soil (Table 4). The Zn concentration in grains ranged from 31.7 mg kg^-1^ (control) to 49.0 and 36.6 mg kg^-1^ under ZnO-NPs and ZnSO_4_ treated soils, respectively. A 17.7% increase in the Zn concentration in grains under ZnO-NPs (T2-T4) treated soil was witnessed as compared to ZnSO_4_ treated soil (T5-T7). The comparison between Zn contents of grains at ZnO-NPs and ZnSO_4_ treated highest doses (T4 and T7) highlighted that Zn content under ZnO-NP treatment was more by 33.9% than ZnSO_4_ treatment (Table 4). It indicated that the Zn use efficiency was significantly higher when Zn was applied in the form ZnO-NPs instead of ZnSO_4_ bulk.

## Discussion

### Zinc nano-fertilization effects on growth and physiology

The advanced nano-engineering may be a handy tool for achieving food security in a sustainable manner. The use of nano-fertilizers improves crop production by enhancing nutrient use efficiency due to reduced losses by more stability and lesser solubility [26, 27]. In the present study, seedling length was 6.8-21.3% higher on the seventh day after putting on petri dishes under ZnO-NPs solution over control (Fig 3 A-C). The mean values were significantly more under ZnO-NPs treatments over the ZnSO_4_ suspension. Our results are in line with the previous research [14]. Wherein, it was noticed that the seedling growth and root and shoot length were increased in lower doses of ZnO-NPs and ZnSO_4_ treatments but negatively affected in concentrations above 50 mg litre^-1^ [14]. Similarly, it was also noticed in earlier work that the lower concentrations of zinc chloride (ZnCl_2_) are beneficial for seedlings growth, whereas, the higher concentration adversely effects vetch root length [43]. Greater stability, reduced solubility and sustained availability of Zn under ZnO-NP treatments might have enhanced the root and shoot length, whereas, under ZnSO_4_ treatments, reduced surface area contact of roots and lower availability might have led to lower root growth [14,43].

The seedling vigor 1 was significantly more (p <0.05) under ZnO-NPs (T2-T3) treatment than that of ZnSO_4_ treatments. It might be due to increased length of seedlings in the treatments T2 and T3 (Table 2). In previous work it has been demonstrated that nanoparticles have potential to augmenting seedling germination and its growth by way of stimulation of enzymatic activities [44–47]. Similarly, it was recorded that the nano technological interventions improves crop health and yield sustainably [48,49]. The hydroxyapatite nano coated urea releases N slowly and uniformly over 60 days, whereas, the traditional bulk fertilizer applied N gets lost within 30-day with uneven release, resulting in reduced N-use efficiency and poor crop growth [50]. Contrarily, some studies have also reported that nano-fertilization adversely affects germination and crop growth. Such type of inconsistency might be due to diverse mode of application of nanoparticles, their shape, size, electronic properties as well as surface coating and species of plants [51,52].

In our experiment, photosynthetic rates of wheat at vegetative stage remained statistically on par across the treatments, which might be due to non-significant difference in chlorophyll content (SPAD values) under different treatments (Table 2). Crop growth and productivity is a function of complex system. The photosynthesis is source of carbon and energy, that the entire system is dependent upon, however, such interactions are not linear and relies on diverse variables [53]. It has been witnessed from several past studies that crop output enhancements are not correlated linearly with gains in photosynthesis [54]. Moreover, in previous work it has also been clearly demonstrated that 15 mg kg^-1^ ferrous and silicon oxide NPs treatment to the soil improved shoot length of maize and barley seedlings by about 20.8% and 8.2%, respectively. Moreover, 25 mg kg^-1^ treatment negatively affected seedlings length of maize which indicates that the crop growth effects due to nanofertilizers is dose dependent [55,56]. Further, our study showed that the tillers plant^-1^ were significantly more under T3 as compared to other treatments. Due to nanofetilization,7.7-24.6% enhancement in tiller count was noticed as compared to T1. The more durability, less leaching and reduced desorption may be the reasons for prolonged nutrient availability to the plants resulting in more tillers (Table 2). Previous reports demonstrated that the plant growth of spinach, cucumber, tomato, wheat and mungbean positively responded to NPs application [57–59]. On the contrary, some reports stated that excessive application of NPs forms reactive oxygen species (ROS) that block nutrient transport channels by their aggregation, which results in reduction in plant growth and quality [60,61].

### Wheat biomass and yield enhancement

The T3 treatment showed significant improvement in biomass and yield. Enhancement in biomass and grain yield under T3 treatment may be due to the more tillers (plant^-1^), grain weight spike^-1^ and spike length (Table 2–3). Previous studies also highlighted that the ZnO-NPs application increased the grain yield of wheat by 56-63% under different doses of ZnO-NPs over the control. However, under higher dose (1,000 mg kg^-1^) significant reduction in yield levels were reported [14]. In another study it has been noticed that ZnO-NPs application increased the number of soybean pods, though, average seed per pod and their size did not change between the treatments [62]. Majority of earlier findings suggested that the conventional fertilizer use efficiency hardly exceeds 30-40% owing to leaching, run off, hydrolysis, drift, evaporation, microbial degradation and photolytic activity [63,64]. Nonetheless, nano-formulation of fertilizers increases their use efficiency due to high stability and higher absorption [28,29].

### Zn partitioning in plant system and biofortification

In our study, Zn concentration in roots, leaves, shoots and grains increased with increase in doses of ZnO-NPs as well as ZnSO_4_. Higher improvement in Zn concentration in grains due to ZnO-NPs (9.5-54.6%) as compared to Zn enrichment owing to ZnSO_4_ (7.6-15.5%) under the current study is in similar lines with earlier findings [60]. However, the partitioning behavior under different plant parts varied significantly. In wheat grain, root and stem, Zn concentrations remained higher with ZnO-NPs application as compared to ZnSO_4_. Contrarily, ZnSO_4_ treatments increased Zn concentration in wheat leaves by 185-270%, whereas under ZnO-NPs the improvement remained lower (54.1-217%). This may be due to enhanced translocation of Zn towards grain due to better source-sink channel and improved use efficiency of applied Zn under ZnO-NPs fertilizer over conventional (bulk) ZnSO_4_ [65]. Our results corroborate previous findings concerned with increased zinc concentration in different plant parts using ZnO-NPs [14].

## Conclusions

The application of nano-engineered fertilizers could be helpful in tackling malnutrition and achieving the food security sustainably by the way of their stimulation in nutrient concentration in edible portion of plants. Fertilizers in nano-scale assure better conservation and management of inputs of plant production as well as protection of environment from pollution. In the present study, it has been demonstrated that zinc oxide nano-particles (ZnO-NPs) are having good potential to improve crop growth and yield as well as Zn concentrations in various plant parts at lower doses over the conventional ZnSO_4_. Present study suggests that 25 mg kg^-1^ ZnO-NPs dose remained most appropriate for maximizing crop growth and yield. Significant enhancement in Zn concentration in edible plant parts owing to ZnO-NPs application would have good implications on tackling Zn hidden hunger and malnutrition of human population. The future research on the similar aspect may focus on evaluating different ZnO-NPs under large plot size in different crops under a cropping system mode. Likewise, a thorough study on understanding Zn release pattern and finding out threshold limits of ZnO-NPs in enhancement in Zn content and yield in different crops could be innovative lines for future work.

## Author Contribution

Conceptualization, A.Y., Y.S.S., R.S.B. and P.K.; Data curation, S.G., S.B., S.Y. and M.K.; Methodology, A.K.C., A.Y. and S.L.M.; Experiment conduction, R.Y., R.R., Abhijeet, A.Y. and S.K.; Data collection and analysis, Shaloo, S.K. and T.S.; Resources, R.Y., R.R., M.S.N. and Abhijeet.; Writing of draft, A.Y., S.G. and R.S.B.; Review and editing: Y.S.S., A.K.C., A.Y.

## Data availability

All relevant raw data about soil and plant are within the paper

## Funding

The study was funded by ICAR-IARI, New Delhi (Project No. CRSC IARISIL 2014028260). The funder had no role in the study design, data collection and analysis, decision to publish, and preparation of the manuscript.

## Competing interests

The authors have declared that no competing interests exist.

## Notes

### Competing Interest Statement

The authors have declared no competing interest.

